# Two contrasting mediodorsal thalamic circuits target the medial prefrontal cortex

**DOI:** 10.1101/2021.01.20.427526

**Authors:** Polina Lyuboslavsky, Alena Kizimenko, Audrey C. Brumback

## Abstract

At the heart of the prefrontal executive and limbic networks is the mediodorsal thalamus (MD). Despite the importance of MD in a broad range of behaviors and neuropsychiatric disorders, virtually nothing is known about the physiology of neurons in MD. Here, we injected the retrograde tracer cholera toxin subunit B (CTB) into the medial prefrontal cortex (mPFC) of adult (8 – 12 week old) male and female wildtype mice. We prepared acute brain slices and used current clamp electrophysiology to measure and compare the intrinsic properties of the neurons in MD that project to mPFC (MD→mPFC neurons). MD→mPFC neurons are located predominantly in the medial (MD-M) and lateral (MD-L) subnuclei of MD. We found that that MD-M→mPFC neurons have longer membrane time constants, higher membrane resistance, less Hyperpolarization and Cyclic Nucleotide gated (HCN) channel activity, and more readily generate action potentials compared to MD-L→mPFC neurons. Additionally, MD-M→mPFC neurons have larger and more complex dendritic arbors compared to MD-L→mPFC neurons. These data demonstrating that the two populations of MD→mPFC neurons have distinct physiologies and morphologies suggests a differential role in thalamocortical information processing and potentially behavior.

## INTRODUCTION

The mediodorsal thalamus (MD) receives inputs from multiple subcortical and cortical brain regions. In particular, MD shares rich reciprocal connections with the prefrontal cortex (PFC). In fact, PFC was originally defined as the cortex that receives projections from MD (Rose and Woolsey, 1948). As part of the prefrontal network, MD participates in cognitive processes such as attention, working memory, behavioral flexibility, and social motivation (Ouhaz et al., 2018). Acquired lesions in MD are associated with executive dysfunction in humans (Hwang et al., 2020). In experimental preparations, intact MD→PFC signaling is required for working memory e.g. (Alexander and Fuster, 1973; Fuster and Alexander, 1973; Parnaudeau et al., 2013; Bolkan et al., 2017), cognitive flexibility (Rikhye et al., 2018), and fear extinction (Lee et al., 2011).

In human neuropsychiatric disorders involving executive dysfunction, there is evidence of abnormal thalamic structure and abnormal connectivity between MD and PFC: autism (Cheon et al., 2011; Maximo and Kana, 2019), schizophrenia (Giraldo-Chica et al., 2018), and epileptic encephalopathy (Warren et al., 2017). Despite the importance of MD in a broad range of behaviors and human disease, virtually nothing is known about the physiology of the neurons in MD.

MD thalamus is composed of three distinct subnuclei (**medial, MD-M**; **central, MD-C**; and **lateral, MD-L**) (Ouhaz et al., 2018)). The majority of MD neurons that project to medial PFC (mPFC) reside in MD-M and MD-L (Alcaraz et al., 2016, 2018; Collins et al., 2018). Connectivity, lesion, and circuit manipulation studies have suggested a generalized model of MD-M playing a role in limbic functions, and MD-L participating in executive functioning (Vertes et al., 2015; Wolff and Vann, 2018; Grodd et al., 2020). Given their distinct patterns of afferents and possible divergent roles in behavior, we hypothesized that MD-M→mPFC and MD-L→mPFC neurons would have differences in intrinsic properties and morphology.

To test this hypothesis, we labeled adult mouse MD→mPFC neurons using retrograde tracers targeted to unilateral prelimbic and infralimbic cortex. In acute brain slices, we used whole-cell patch clamp recordings and morphological reconstruction to analyze and compare subthreshold and suprathreshold intrinsic properties and dendritic morphology of labeled MD-M→mPFC and MD-L→mPFC neurons. We found significant differences in neuronal morphology, intrinsic membrane physiology, and action potential dynamics between MD-L and MD-M neurons. These findings suggest that MD provides at least two ascending information pathways to mPFC which process physiological information in fundamentally different ways.

## MATERIALS AND METHODS

### Animals

All experiments were conducted in accordance with procedures established by the Institutional Animal Care and Use Committee (IACUC) at The University of Texas at Austin. Most mice were bred in-house and weaned at 3 weeks of age. Adult mice were group-housed by sex in open-top cages and fed *ad libitum*. Animals were kept in reverse lighting conditions (12 h / 12 h dark / light cycle) and were typically euthanized for patch clamp physiology experiments in the mornings, around the beginning of the animals’ dark cycle. In our experiments, we used male and female C57Bl/6J mice from Jackson Laboratories (strain # 000664), *Fmr1* WT mice (a gift from Dan Johnston (Huber et al., 2002)), and *Drd2-Cre* mice (B6.FVB(Cg)-Tg(Drd2-cre)ER44Gsat/Mmcd; MMRRC strain 032108). There were no systematic differences in electrophysiological properties or cellular morphology between male and female mice, or between the various strains of wildtype mice used.

### Fluorescent labeling of specific neuron populations

Mice were anesthetized with 2% isoflurane and mounted in a stereotactic frame. Craniotomies were made according to stereotaxic coordinates relative to Bregma. To label mediodorsal thalamus (MD) neurons that project to ipsilateral prelimbic and infralimbic cortices (ipsilateral mPFC), we injected fluorescently-labeled cholera toxin subunit B (CTB, 500 µg / 100 µL, Molecular Probes, Thermo Fisher Scientific, Waltham, MA, USA) into ipsilateral mPFC (Nanofil Syringe and Pump UMP3, World Precision Instruments). Coordinates for injection into ipsilateral mPFC were (in mm relative to Bregma): –1.7 anterior–posterior (AP), +0.3 mediolateral (ML) and –2.75 dorsoventral (DV). After needle insertion, we waited 5 minutes before starting the injection of 450 nL at 100 nL / minute into mPFC. We waited 5 minutes after the end of the injection before slowly withdrawing the syringe needle. We waited 3–6 days following retrograde tracer injections before performing experiments. At the time of the experiments, we visually verified that retrograde tracer injections were targeted appropriately, and that tracer was not present in nearby structures.

### Slice preparation

Except as noted, all reagents for patch clamp electrophysiology solutions were purchased from Sigma Aldrich. We used ice-cold cutting solution containing (in mM): 205 sucrose, 25 NaHCO_3_, 2.5 KCl, 1.25 NaH_2_PO_4_, 7 MgCl_2_, 7 dextrose, 3 Na pyruvate, 1.3 sodium ascorbate, and 0.5 CaCl_2_ bubbled with 95% O_2_ / 5% CO_2_. Mice 8 to 12 weeks old were deeply anesthetized with intraperitoneal injection of ketamine / xylazine (200/100 mg/kg; Acor/Dechra), transcardially perfused with 20 mL of ice-cold cutting solution, and then decapitated. We prepared 300 µm coronal slices (Leica VT1200). Slices were incubated in holding solution containing (in mM): 125 NaCl, 25 NaHCO_3_, 2.5 KCl, 1.25 NaH_2_PO_4_, 25 dextrose, 2 CaCl_2_, 2 MgCl_2_, 1.3 sodium ascorbate, and 3 Na pyruvate at 37±1°C for 30 minutes. Slices were then kept for at least 30 minutes at room temperature before being used for recordings.

### Intracellular recordings

Artificial cerebrospinal fluid (ACSF) contained (in mM): 125 NaCl, 25 NaHCO_3_, 12.5 dextrose, 2.5 KCl, 1.25 NaH_2_PO_4_ 2 CaCl_2_ and 1 MgCl_2_. Slices were continuously perfused with ACSF in an immersion chamber (Warner Instruments) with temperature maintained at 32.5 ± 1°C (Warner Instruments TC-324C). We did not add synaptic blockers to the ACSF.

Somatic whole-cell patch recordings were obtained from visually-identified retrogradely-labeled neurons in the medial (MD-M) or lateral (MD-L) subnuclei, using DODT (Zen 2.5 blue addition, Zen pro) contrast microscopy and epifluorescence on an upright microscope (Zeiss Examiner D1). Patch electrodes (tip resistance = 3–6 MΩ) were filled with the following (in mM): 118 K-gluconate, 10 KCl, 10 HEPES, 4 MgATP, 1 EGTA, 0.3 Na3GTP and 0.3% biocytin (pH adjusted to 7.2 with KOH; 282 mOsM). For some cells, we added 16 µM Alexa 488 or Alexa 594 to to visualize the dendritic arbor under epifluorescence. There was no qualitative difference in the recordings made from cells with Alexa dye in the internal solution and those without. Recordings were made with Clampex 10.7 software running a Multiclamp 700B (Molecular Devices). Signals were digitized at 20,000 Hz and lowpass filtered at 4,000 Hz.

Gigaohm seal was achieved in voltage clamp, and after establishing whole-cell configuration, we immediately switched to current clamp to measure the resting membrane potential. After the membrane potential stabilized, data were collected at the cell’s resting membrane potential. Then, all cells were provided steady state current to maintain the membrane potential at −65 +/- 3 mV. With the exception of the RMP measurement, all data reported here are taken from the recordings performed at −65 mV. Series resistance was usually 10–20 MΩ, and experiments were discontinued above 30 MΩ or if action potentials failed to overshoot 0 mV. Experiments typically lasted <30 minutes total. Membrane potentials are reported without correcting for liquid membrane junction potential.

### Morphology

After recording, slices were stored overnight at 4°C in 4% PFA (Electron Microscopy science) and moved to phosphate-buffered saline (PBS) the following morning. Within 60 days, we stained cells with DAB to reveal biotin filling. First, we washed slices in 0.1M PBS for 30 minutes (6 x 5 minutes) with the consecutive washing in 1% Triton (Fisher Chemical) and 0.5% H_2_O_2_ (Fisher Chemical) solutions in PBS for 30 minutes each at room temperature (RT). After washing in 0.1M PBS again for 30 minutes, we incubated slices in Reagent A + Reagent B + 0.5% TritonX + 0.1% BSA (Sigma-Aldrich) in PB for 2 days at 4^°C^ (ABC reagent, VectaStain Elite ABC-HRP kit, Vector laboratories). After the 2 day incubation, we washed slices in PBS for an hour (6 x 10 minutes). Then we incubated slices with DAB (DAB substrate kit, Vector laboratories) solution for one hour at RT. After that we added 3% H_2_O_2_ to the slices (one drop every 20 minutes) until they reached the desirable brown color. Then we washed slices again for 30 minutes and dehydrated them in ascending concentrations of glycerol (Fisher Chemical) from 20% to 100% for 10 minutes per concentration. We mounted slices on slides using 100% glycerol as the mounting medium. We reconstructed cells using the Neurolucida System (MBF Bioscience Inc. Williston, VT) on Leitz Diaplan microscope and MicroFire Optronics camera using a 40x objective. We placed the zero point in the center of the soma (at the widest part of the soma) and traced dendrites using a fine focusing wheel. We considered branches as cut when they visibly went out of the slice. We considered cells usable for analysis if they had fully-visualized dendritic arbors with well-defined terminals and with the majority of branches being present. All neuronal morphologies will be cataloged at www.NeuroMorpho.org.

### Histology

To create an atlas of fixed brain slice images onto which we could map all recorded cells, we injected an 11 week old C57Bl/6 wildtype mouse with CTB into the prelimbic and infralimbic cortices as above. The animal was deeply anesthetized with ketamine/xylazine and transcardially perfused with paraformaldehyde (PFA (Sigma-Aldrich)) 4% in phosphate-buffered saline (1x PBS). We prepared 150 µm thick coronal slices with DAPI-containing mounting medium (VectaShield HardSet with DAPI, Vector laboratories). We imaged the slices at 5x using Zeiss Axio Imager 2 and stitched images together using AxioVision software.

### Data analysis: Electrophysiology

We estimated the resting membrane potential (RMP) as the membrane voltage recorded immediately after switching to current clamp configuration. We calculated membrane time constant (T_m_) as the time at which the membrane voltage decayed to 1/e of the initial value following 1000 ms x –10 pA current steps averaged over 10 repetitions. We estimated membrane resistance (R_m_) from the steady-state voltage change measured in response to 1000 ms current steps ranging from −60 to +60 pA in 5 pA increments. We calculated R_m_ as the slope of the linear relationship between subthreshold steady-state voltage and input current.

To estimate HCN channel activity, we calculated the voltage sag in response to 1000 ms current steps ranging from −250 to −150 pA in 25 pA increments. We identified the sweep that elicited a peak hyperpolarization of −100 mV and measured voltage sag (minimum voltage) – (steady state voltage).

We measured the absolute value of the afterhyperpolarization (AHP) following 1000 ms depolarizing current steps as the minimum voltage in the 250 ms following the offset of the current step. To measure the net AHP compared to baseline, we subtracted the AHP absolute value from the membrane potential 4 seconds following the offset of the current step (when the membrane potential had returned to baseline). For experiments in which we washed on ZD7288 (20 μM; Tocris Cat#1000) to block HCN channel activity, we measured the membrane potential in the post-drug state at the same time point at which the AHP was measured in the pre-drug state.

We quantified action potential firing during one second current steps from 0 to +60 pA in 5 pA increments and from 0 to +350 pA in 25 pA increments. We estimated the action potential threshold as the point at which the third derivative of the membrane potential was maximal. We categorized action potentials (“spikes”) as belonging to the following categories: rebound, burst, tonic, or total. We counted spikes as belonging to the rebound if they occurred within 1 second after the offset of a negative current step. We classified spikes as belonging to a depolarization-induced burst if they occurred within 500 ms of the onset of a depolarizing current step, had an inter-spike interval of < 40 milliseconds, had an inter-ISI ratio of 0.50 − 1.99, and the total number of spikes in that sweep’s step was less than or equal to 15. During depolarizing current steps containing 15 or fewer total spikes, the first spike to have an inter-ISI ratio of ≥ 2 was classified as the first tonic spike. The rest of the spikes in the train following the first tonic spike were also classified as tonic. In current steps in which the number of spikes during the step exceeded 15, all spikes were grouped together as “total spikes”, as we could not reliably distinguish burst from tonic spiking. To quantify accommodation, we identified the lowest amplitude depolarizing sweep to have ≥12 tonic action potentials during the depolarizing step. We estimated the accommodation index of tonic spikes by calculating the slope of the linear relationship between the interspike intervals for each successive action potential during the current step (**Figure 5**).

### Data analysis: Neuronal morphology

We used Neurolucida Explorer (MBF Bioscience Inc. Williston, VT) to analyze the cell’s morphological parameters. We measured dendritic length (sum total length of all branches of the neuron (µm)) and average dendritic length: dendritic length / number of primary dendrites. We assessed total number of branching points (nodes) per cell and total number of last order branches (terminals) per cell. In addition to first order branches, we also analyzed the number of secondary and tertiary branches. We used convex hull analysis to measure area (µm^2^) and volume (µm^3^). These parameters correspond to the area and volume of a convex polygon that connects tips of distal processes. For Sholl analysis (Sholl, 1953), we placed concentric circles in 15 µm radius increments around the cell centered at the cell body. We counted the number of points where cell processes intersect with Sholl circles. Within each circle, we also measured dendritic length. We calculated cellular complexity using the following formula:

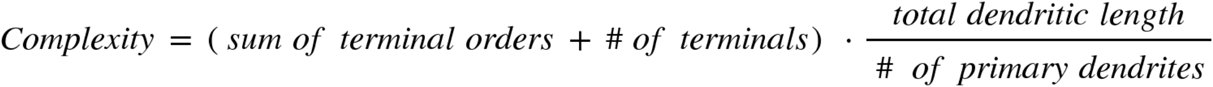

For each terminal in a cell, the *terminal order* is the total number of sister branches encountered as one traces a path from that terminal to the cell body. The *sum of terminal orders* is the sum of all *terminal order*s for each cell.

### Statistics

We used the ‘sampsizepwr’ function in MATLAB to calculate sample sizes based on preliminary data. For patch clamp electrophysiology experiments, we estimated that between MD-M and MD-L, to detect a difference in membrane time constant of 25% with a standard deviation of 20 ms, given α of 0.05 and power of 0.8, we required at least 16 cells per group. For neuronal morphology experiments, we estimated that between MD-M and MD-L, to detect a difference in complexity of 100% with a standard deviation of 50,000 μm, given α of 0.05 and power of 0.8, we required 5 cells per group.

For all data sets, we tested if data followed a Gaussian distribution. If data were normally distributed, we used two-tailed unpaired Student’s t-tests to compare two groups or two-tailed paired Student’s t-tests for comparison of predrug / postdrug treatments in individual neurons. For data that were not normally distributed, we used the non-parametric Mann-Whitney test. To test significance of differences between groups in action potential firing as a function of input current and for Sholl analyses, we used two-way ANOVA with Sidak’s test to correct for multiple comparisons.

All electrophysiology quantifications were performed using custom-written code in Matlab (to be made publicly available on GitHub). All statistical analyses except for power analysis were performed using GraphPad Prism version 8.0.0 (GraphPad Software, San Diego, California USA, www.graphpad.com). All graphs were made using GraphPad Prism and transferred to Adobe Illustrator version 24.3 for final presentation.

All sample sizes are reported as the number of neurons. We typically prepared 3 brain slices per animal, and recorded 1 neuron per brain slice. In the text, all data are reported as mean ± standard deviation. In the figures, all data are plotted as mean ± standard error. We defined a ≤ 0.05 with * p < 0.05; ** p < 0.01; *** p < 0.001; **** p < 0.0001.

We removed one outlier from the rebound action potential data set (rheobase = −60 mV) based on GraphPad Prism’s ROUT method of identifying outliers (Q = 1%; Motulsky and Brown 2006).

## RESULTS

To test the hypothesis that the MD→mPFC projection consists of different physiological and morphological cell types, we used whole cell patch clamp electrophysiology coupled with analysis of cellular morphology. We injected a fluorescent retrograde tracer (CTB) into ipsilateral prelimbic and infralimbic cortices (**Figure 1A,B**). After 3-6 days, we euthanized the animal, prepared acute coronal brain slices, and recorded from individual visually-identified MD→mPFC neurons. We distinguished the medial (MD-M), central (MD-C), and lateral (MD-L) subnuclei within MD based on the distribution of fluorescent labeling and distance from midline. The vast majority of labeled neurons were present in the medial (MD-M) and lateral (MD-L) divisions (**Figure 1C-E**).

**Figure 1:**
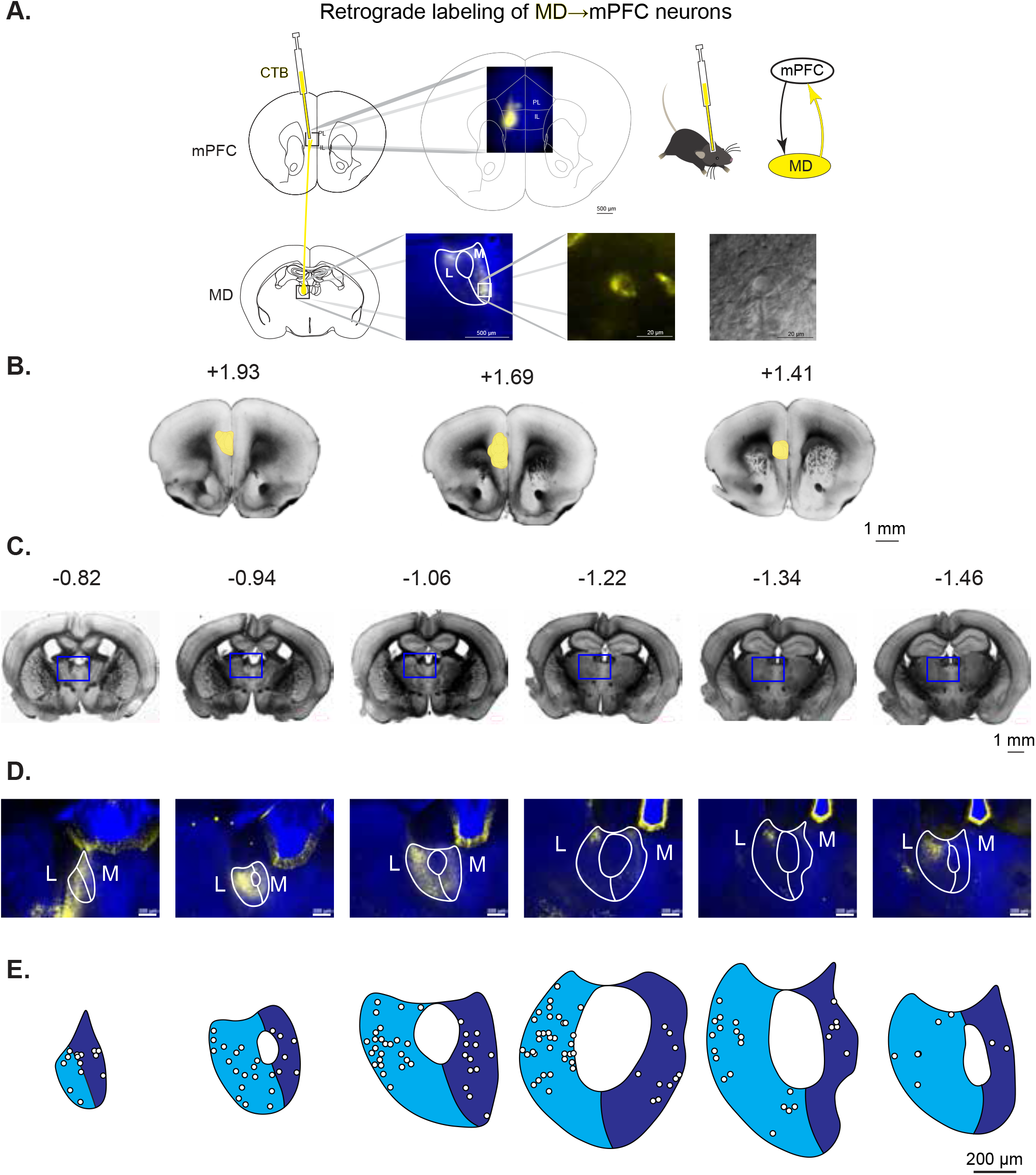
MD→mPFC neurons predominantly reside in MD-M and MD-L. **A**. Adult wildtype male and female mice were stereotaxically injected with the retrograde fluorescent tracer CTB into the right medial prefrontal cortex. Fluorescently-labeled MD→mPFC neurons were visually identified for patch clamp electrophysiology experiments. **B**. Bright field photomicrographs of coronal mouse brain slices with the estimated distance from Bregma (in mm) demonstrating the locations of the CTB injection sites in mPFC. **C**. Bright field photomicrographs of coronal mouse brain slices with the estimated distance from Bregma (in mm) demonstrating labeled cell bodies of MD→mPFC neurons. **D**. Photomicrographs (10x) of the boxed areas from **C** demonstrating MD→mPFC labeled neurons (yellow) in each of the representative slices. **E**. Map of the approximate locations of all recorded neurons reported in this manuscript placed on a single representative atlas drawing for each coronal slice. Medial MD (MD-M) is dark blue. Lateral MD (MD-L) is highlighted in cyan. This color scheme is used throughout the paper to denote MD-M→mPFC and MD-L→mPFC neurons.

### Subthreshold properties: membrane time constant and input resistance

We compared the intrinsic membrane properties of medial and lateral MD→mPFC neurons. There was no significant difference in resting membrane potential (**Figure 2B**; MD-M: −59.29 ± 8.85 mV (n = 42 neurons); MD-L: −59.28 ± 7.79 (n = 110 neurons); Mann-Whitney test; *U* = 2154; *p* = 0.52).

**Figure 2:**
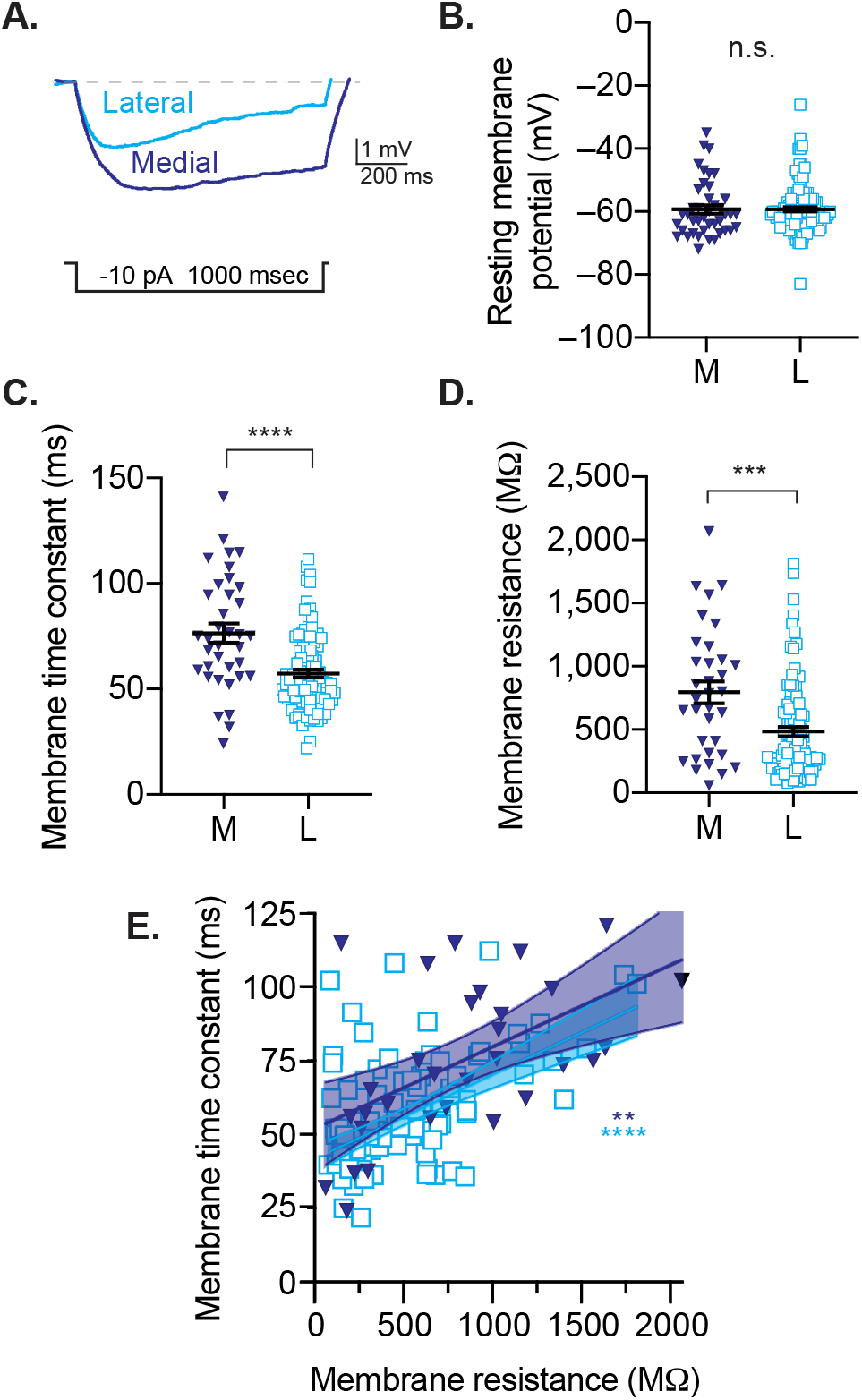
Subthreshold membrane properties of prefrontal medial (MD-M) and lateral (MD-L) thalamocortical neurons. **A**. Estimates of membrane time constant and input resistance using low-amplitude hyper-polarizing steps. Neurons were held in current clamp at −65 mV. **B**. There was no difference in resting membrane potentials between groups. **C**. Lateral neurons have faster membrane times constant compared to medial neurons. **D**. Membrane resistance is decreased in lateral neurons compared to medial neurons. **E**. For both medial and lateral neurons, membrane time constant is proportional to membrane resistance. Scatter plots show all data points with the mean ± SEM. Significance is indicated as not significant (n.s.; p>0.05), or significant at p<0.01 (**), p<0.001 (***), p<0.0001 (****).

The membrane time constant influences a neuron’s intrinsic and synaptic physiology. We estimated the membrane time constant with repeated small-amplitude (−10 pA) current steps (**Figure 2A**). We found that lateral neurons had shorter membrane time constants than medial neurons (**Figure 2C**; MD-M: 76.46 ± 26.92 ms (n = 36 neurons); MD-L: 57.33 ± 18.38 ms (n = 101 neurons); Mann-Whitney test; *U* = 993.5; *p* = <0.0001). Membrane time constant is the product of cellular membrane resistance *R*_*m*_ and cellular capacitance *C*_*m*_. We estimated membrane resistance based on the voltage response to subthreshold current steps, and found that lateral neurons had lower membrane resistance than medial cells (**Figure 2D**; MD-M: 795 ± 502.6 (n = 34 neurons); MD-L: 484.9 ± 372.7 (n = 101 neurons); Mann-Whitney test; *U* = 1056; *p* = 0.0007). We found that in both medial and lateral cells, membrane time constant correlated with membrane resistance (**Figure 2E**; 95% CI for slope of linear regression: MD-M: 0.012 to 0.043; MD-L: 0.021 to 0.036; Spearman correlation: MD-M: *r*_*s*_ = 0.493; *p* = 0.0036; *r*_*s*_ = MD-L 0.432; *p* < 0.0001).

### Subthreshold properties: HCN channel related properties

*R*_*m*_ depends on multiple factors including Hyperpolarization and Cyclic Nucleotide (HCN) activated channels. We estimated HCN channel activity by measuring the voltage sag in response to hyperpolarizing current steps (**Figure 3A**). We found that lateral cells had a higher degree of voltage sag than medial cells (**Figure 3B**; MD-M: 6.398 ± 3.957 (n = 43 neurons); MD-L: 8.627 ± 4.521 (n = 109 neurons); Mann-Whitney test; *U* = 1683; *p* = 0.0066). To estimate the relationship between cellular membrane resistance and the HCN channel activity, we calculated the relationship between *R*_*m*_ and the voltage sag for each neuron. We found that membrane resistance was inversely related to voltage sag (**Figure 3C**; 95% CI for slope of linear regression: MD-M: −106.5 to −44.19; MD-L: −62.07 to −38.63; Spearman correlation: MD-M: *r*_*s*_ = −0.6731; *p* <0.0001; MD-L: *r*_*s*_ = −0.654; *p* < 0.0001). Voltage sag was abolished in both cell types following bath application of ZD7288, an inhibitor of HCN channel activity (**Figure 3D**; MD-M: pre-ZD: 4.02 ± 1.783; post-ZD: 0.2996 ± 0.6083 mV (n = 23 neurons; Two-tailed paired T test; *t* = 6.770, *df* = 9; p <0.0001); MD-L: pre-ZD: 8.726 ± 3.169; post-ZD: 0.4577 ± 0.4892 mV (n = 18 neurons); Two-tailed paired T test; *t* = 11.15, *df* = 17; p <0.0001). Application of ZD7288 to MD-L cells increased *R*_*m*_ to MD-M levels (**Figure 3E**; pre-ZD: 476.7 ± 349.4 MΩ; post-ZD: 903.8 ± 389.4 MΩ (*n* = 17 neurons); Wilcoxon sum of ranks 149, −4, *W* = 145; p = 0.0001). There was no effect of ZD7288 on *R*_*m*_ in MD-M neurons (pre-ZD: 854.4 ± 411.5 MΩ; post-ZD: 876.3 ± 467.5 MΩ (*n* = 10 neurons); *t* = 0.1481, *df* = 9; *p* = 0.8855).

**Figure 3:**
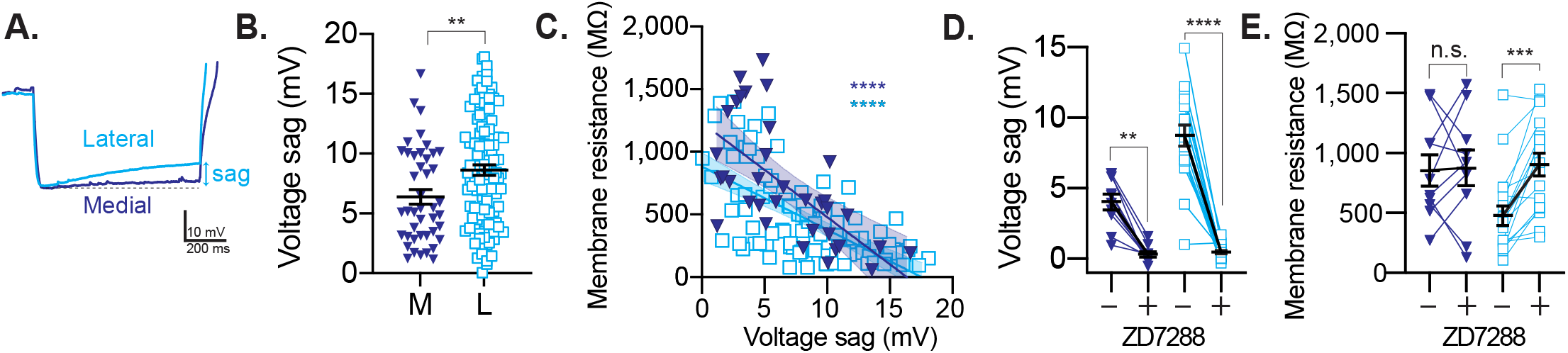
MD-L→mPFC neurons have increased voltage sag compared to MD-M→mPFC neurons. **A**. Representative voltage traces of MD-M and MD-L neurons in response to negative current steps causing hyperpolarization to −100 mV. Rebound action potentials are truncated to simplify display. **B**. MD-L cells have increased voltage sag compared to MD-M neurons. **C**. Neurons with higher voltage sag have lower membrane resistance. **D**. In both cell types, voltage sag decreases following bath application of the HCN channel inhibitor ZD7288 (20 µM). **E**. In MD-L neurons, bath application of ZD7288 increases membrane resistance to the MD-M range. Scatter plots show all data points with the mean ± SEM. Significance is indicated as not significant (n.s.), p<0.05 (*), p<0.01 (**), p<0.001 (***), p<0.0001 (***).

Following a depolarizing current step, the membrane potential hyperpolarizes transiently before returning to baseline (**Figure 4A**). This afterhyperpolarization (AHP) is abolished by ZD7288, an inhibitor of HCN channel activity (**Figure 4D**), and thus we used AHP amplitude as a complementary measure to voltage sag to estimate HCN channel activity in MD→mPFC neurons (Fan et al., 2005). We measured the amplitude of the AHP as the most negative potential in the 500 ms after the offset of the most positive (+350 pA) current step (at which there is no difference between groups in the number of action potentials generated). Lateral cells had a more robust AHP than medial cells (**Figure 4B**; MD-M: −73.18 ± 7.71 mV (n = 40 neurons); MD-L: −76.87 ± 6.85 mV (n = 103 neurons); Two-tailed T test, t=2.797, df=141; p = 0.0059). For both cell types, the AHP correlated with the voltage sag (**Figure 4C**; 95% CI for slope of linear regression: MD-M: −1.858 to −0.8596; MD-L: −1.145 to −0.6880; Spearman correlation: MD-M: *r*_*s*_ = −0.6716; p < 0.0001; MD-L: *r*_*s*_ = −0.613; p < 0.0001). The AHP was abolished in the presence of ZD7288, a selective blocker of HCN channels (**Figure 4D**; pre-ZD: −12.43 ± 6.179 (n = 25 neurons); post-ZD: 6.185 ± 3.51 (n = 23 neurons); Two-tailed paired T test; *t* = 11.91, *df* = 22; p <0.0001).

**Figure 4.**
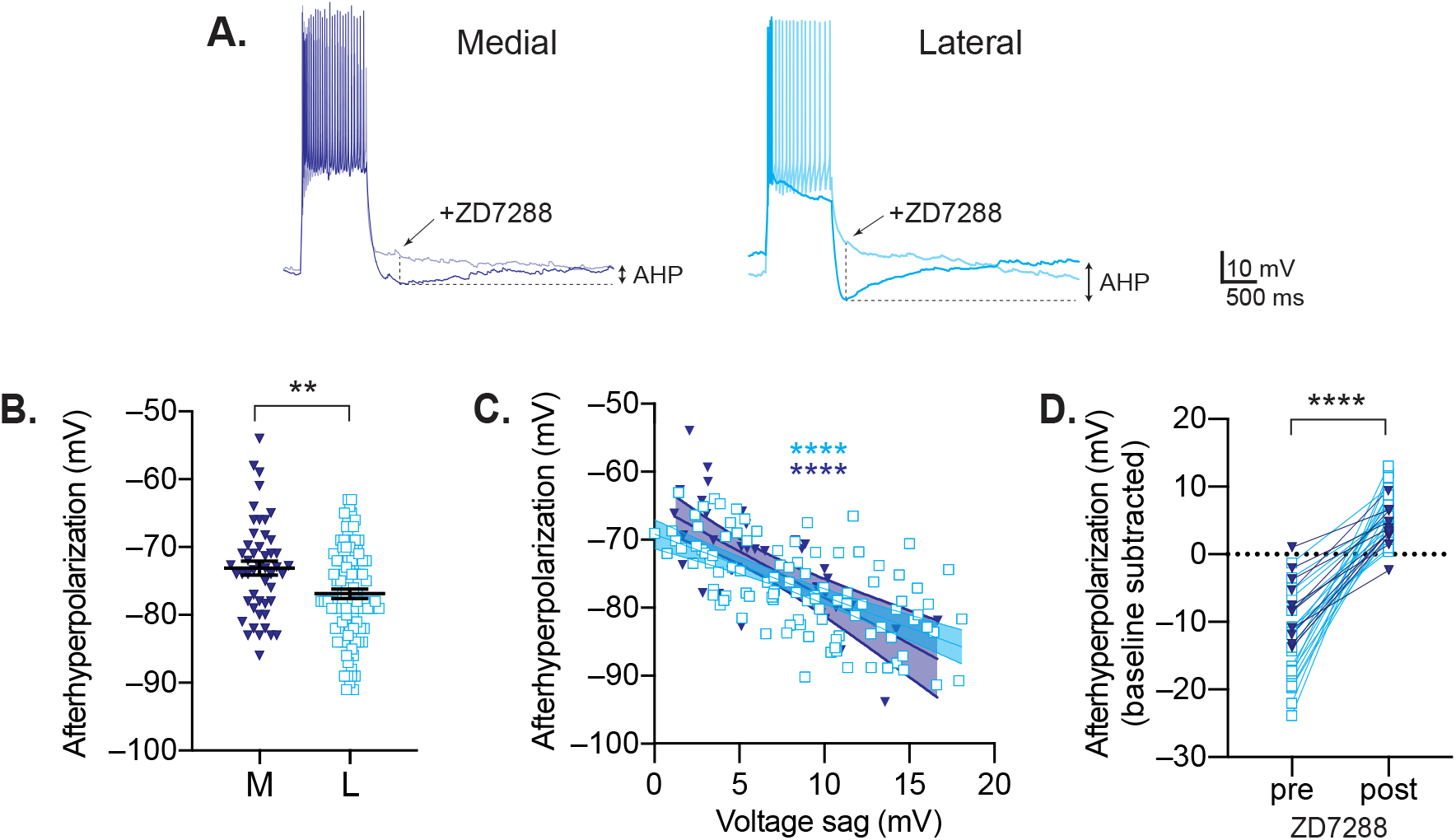
Subregion differences in HCN-channel-dependent afterhyperpolarization (AHP). **A**. Current clamp recordings of medial and lateral MD→mPFC neurons in response to a +100 pA depolarizing current step pre (dark lines) and post (light lines) bath application of ZD7288 (20 μM), a selective inhibitor of HCN channels. **B**. In control conditions, AHP amplitude is greater in lateral cells compared to medial cells. **C**. AHP amplitude is correlated with another measure of HCN-channel activity, voltage sag. **D**. The AHP is abolished with bath application of ZD7288 in both MD-M→mPFC and MD-L→mPFC neurons. Scatter plots show all data points with the mean ± SEM. Significance is indicated as p<0.01 (**), p<0.0001 (****).

**Figure 5.**
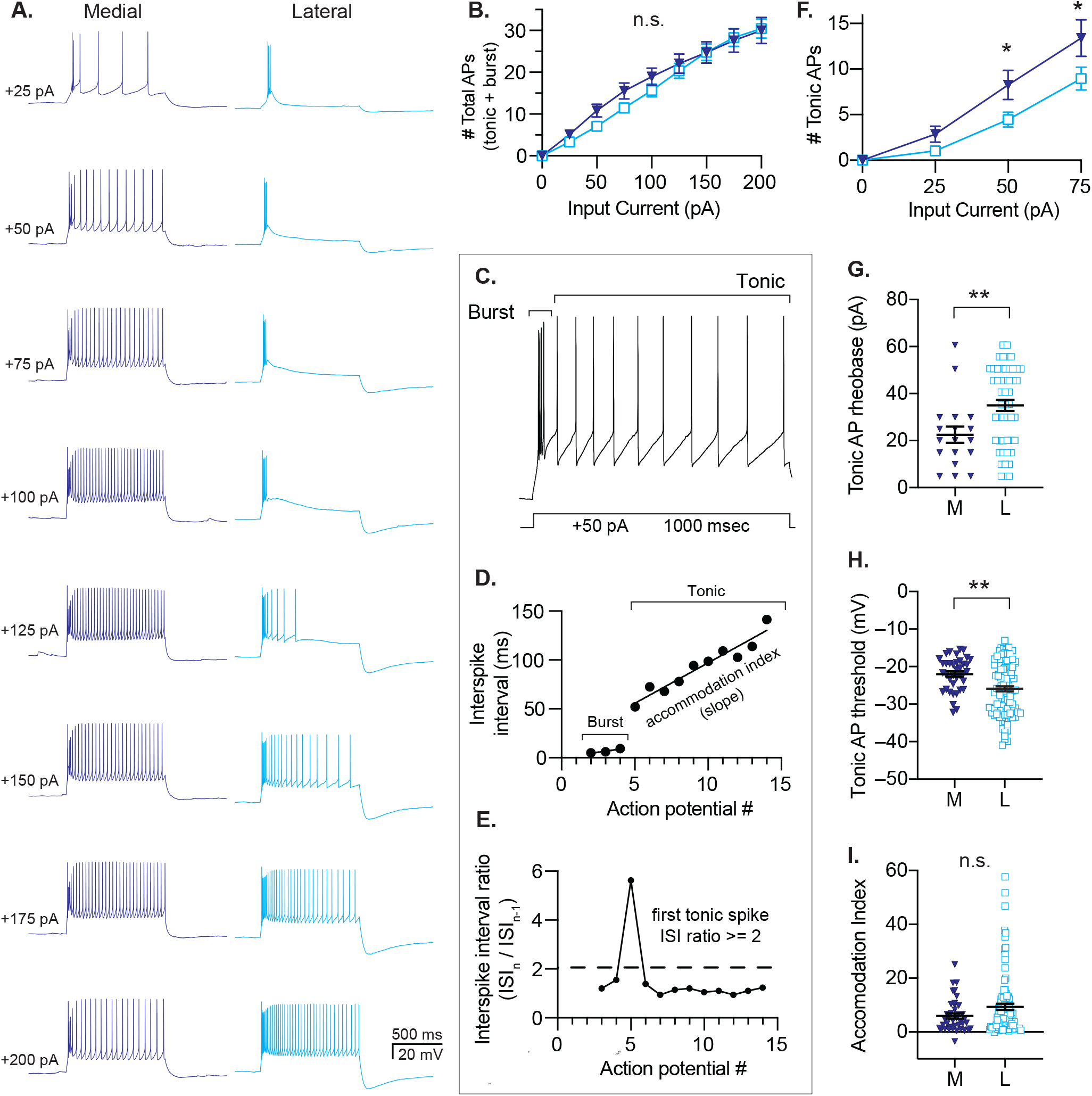
Action potential generation differs between lateral and medial MD→mPFC neurons. **A**. Representative voltage responses to 1000 ms current steps of increasing amplitude. **B**. Lateral cells fire similar numbers of action potentials in response to depolarizing current steps compared to medial cells. **C-E**. Action potentials are classified as “burst” or “tonic”. **C**. Representative voltage response to a +50 pA current step demonstrating the two phases of action potential firing. **D**. Burst spikes occur at the beginning of the current step, cluster as 2-8 spikes with interspike intervals of <40 ms. **E**. The first tonic spike is identified when there is a sharp rise in the ISI interval (>100% increase = ISI ratio >2). See Methods for details. **F**. Lateral MD**→**mPFC neurons generate fewer tonic action potentials in response to depolarizing current steps. **G**.The rheobase for tonic spike generation is increased in lateral neurons. **H**. Action potential threshold is more depolarized in medial cells compared to lateral cells. **I**. Accomodation index for tonic spikes is the slope of the linear fit of ISI vs. action potential # (as in **D**). Both groups had cells with a broad range of accomodation indices. There was no significant difference in mean accomodation index between groups. Scatter plots show all data points with the mean ± SEM. Significance is indicated as not significant (n.s.; p>0.05), or significant at p<0.05 (*), p<0.01 (**) **B, F**: Sidak’s test. **G, H, I**: Mann-Whitney, two tailed.

### Tonic spiking properties

We measured the properties of action potential spiking in response to depolarizing currents steps (**Figure 5A**). Thalamic neurons engage in two modes of action potential firing: bursting and tonic firing (**Figure 5C-E**). Burst spikes are produced by low-voltage activated calcium channels (Ca_V_3, previously known as T-type) along with voltage-gated sodium and potassium channels, whereas tonic spikes do not require calcium channels.

Because they are mediated by different mechanisms, we analyzed burst and tonic spikes separately (**Figure 5C-E**). We found that compared to lateral cells, medial neurons had an increased input-output curve for tonic spiking (**Figure 5F**; Two-way ANOVA: MD subnucleus *F* (1, 592) = 11.17; *p* = 0.0009; Sidak’s test: 50 − 75 mV, adjusted *p* < 0.05). Accordingly, medial neurons had a lower rheobase (the minimum current required to elicit spiking) for tonic spiking compared to lateral neurons (**Figure 5G**; MD-M: 22.47 ± 14.77 pA (n = 18 neurons); MD-L: 34.95 ± 16.79 pA (n = 50 neurons); Mann-Whitney test; *U* = 266; *p* = 0.0095). In contrast, the membrane potential threshold for tonic spiking was higher in medial cells compared to lateral cells (**Figure 5F**; MD-M: −21.8 ± 4.481 mV (*n* = 38 neurons); MD-L: −25.98 ± 6.877 mV (*n* = 101 neurons); Mann-Whitney test; *U* = 1235; *p* = 0.0011). When provided with long-lasting inputs, neurons tend to fire action potentials that are regularly spaced (non-accommodating) or with increasing inter-spike intervals (accommodating). Both cell populations had accommodating and non-accommodating neurons and there was no difference in the mean between the groups (**Figure 5I**; MD-M: 6 ± 6.09 (n = 39 neurons); MD-L: 9.35 ± 11.25 (n = 101 neurons); Mann-Whitney test; *U* = 1672; p = 0.1681).

### Burst spiking properties

In thalamic neurons, bursts of 2 or more tightly-spaced action potentials can be elicited in response to direct depolarization (**Figure 5C** & **6A**). Lateral MD→mPFC neurons had a higher rheobase for firing bursts compared to medial neurons (**Figure 6B**; MD-M: 9.09 ± 5.55 pA (n = 31 neurons); MD-L: 15.99 ± 9.99 pA (n = 95 neurons); Mann-Whitney test; *U* = 820; *p* = 0.0002). Similar to differences in tonic spike threshold, we identified a more hyperpolarized action potential threshold in the lateral neurons compared to medial neurons with burst firing (**Figure 6C**. MD-M: −30.43 ± 5.528 mV (n = 42 neurons); MD-L: −33.76 ± 7.435 mV (n = 109 neurons); Mann-Whitney test; *U* = 1626; *p* = 0.0056). The overall input / output curve for burst spikes was unchanged between the two groups (**Figure 6D**. Two-way ANOVA, *p* = 0.1322)

**Figure 6.**
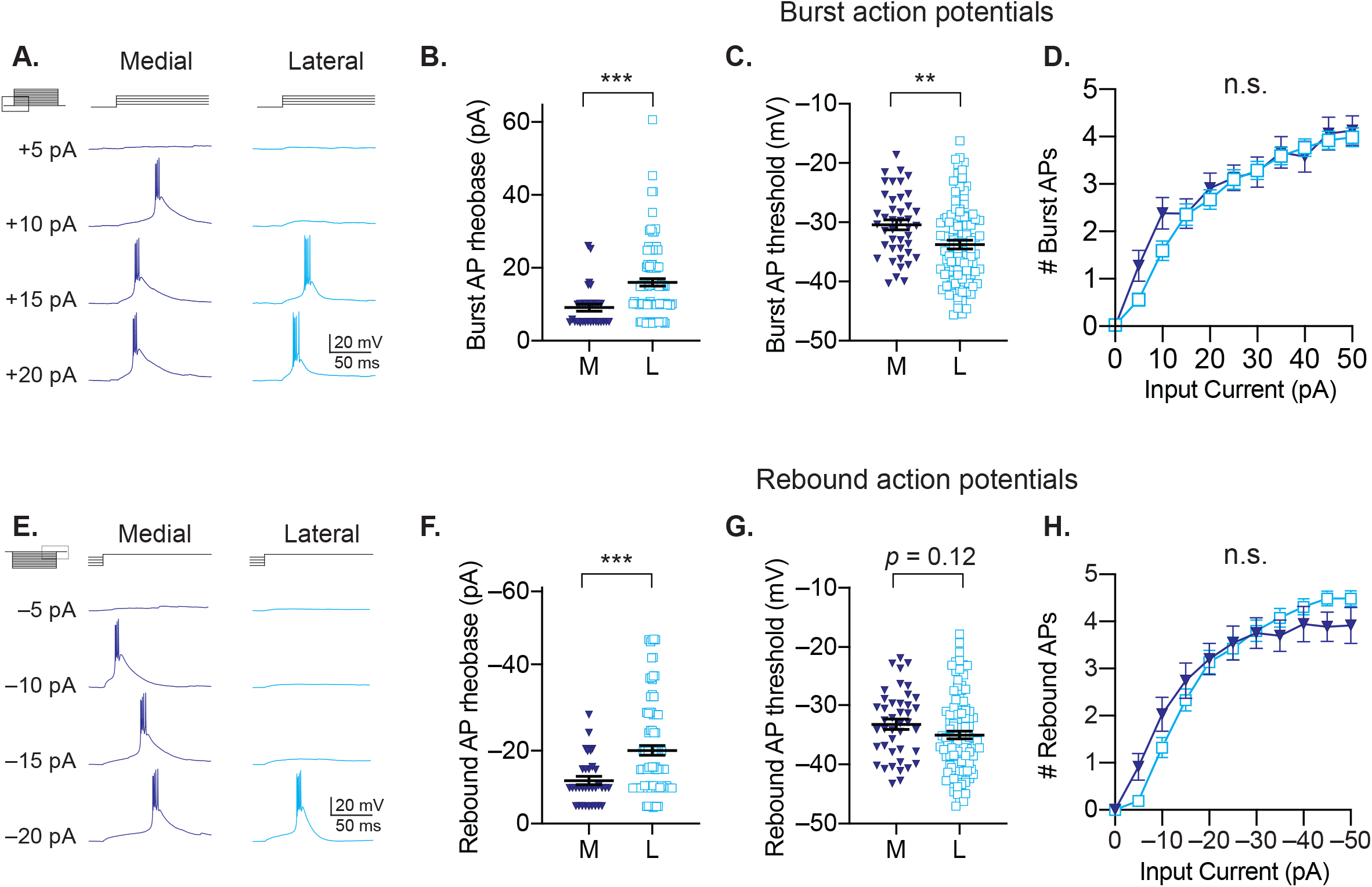
Examination of burst spikes in response to depolarization (*top*) and rebound burst spikes following membrane hyperpolarization (*bottom*). **A**. Examples of voltage responses of medial and lateral MD**→**mPFC neurons to depolarizing current steps. **B**. Lateral cells require more input current to fire burst action potentials (rheobase) in response to depolarizing current steps. **C**. Lateral cells have a more hyperpolarized action potential threshold compared to medial cells for bursts evoked by depolarizing current steps. **D**. The input/output curves for burst firing are not different between groups. **E**. Examples of voltage responses of medial and lateral MD**→**mPFC neurons following hyperpolarizing current steps of varying amplitude. **F**. Lateral cells require more input current to fire rebound action potentials (rheobase) in response to hyperpolarizing current steps. **G**. Lateral cells have a more hyperpolarized action potential threshold compared to medial cells for bursts evoked by hyperpolarizing current steps. **H**. The input/output curves are not different between groups. Scatter plots show all data points with the mean ± SEM. Significance is indicated as not significant (n.s.; p>0.05), or significant at p<0.01 (**) or p<0.001 (***). For **B, C, F, G**: two-tailed Mann-Whitney test. For **D, H**: Sidak’s test.

Bursting can also be induced in response to release of hyperpolarization, so we tested the rheobase for “rebound” burst spikes following hyperpolarizing steps of varying amplitude (**Figure 6E**). Lateral cells required more hyperpolarization to induce rebound bursting compared to medial cells (**Figure 6F**; MD-M: −11.83 ± 6.52 mV (n = 34 neurons); MD-L: −20.09 ± 12.6 mV (n = 93 neurons); Mann-Whitney test; *U* = 907.5; *p* = 0.0002). The difference in rebound action potential threshold between groups did not reach statistical significance (**Figure 6G**; MD-M: − 33.22 ± 5.687 mV (n = 42 neurons); MD-L: −35.04 ± 6.674 mV (n = 106 neurons); Two-tailed t-test; *t* = 1.552, *df* = 146; *p* = 0.1227). The overall input / output curve for rebound burst spikes was also similar between the two groups (**Figure 6H**. Two-way ANOVA, *p* = 0.9879).

### Cellular Morphology

We filled cells with biocytin and processed for histological reconstruction using Neurolucida. **Figure 7A** shows the approximate locations of the reconstructed neurons. Sholl analysis (**Figure 7C**) demonstrated that, in general, MD-M neurons were larger and more complex than neurons from MD-L. Sholl length (dendrite length within each concentric circle) was greater for medial neurons than lateral neurons (**Figure 7F**; Two-way ANOVA, main effect of MD subnucleus; *F*(1, 315) = 73.22; *p* < 0.0001; Sidak’s multiple comparisons test: 90 − 165 μm from the soma, adjusted *p* values < 0.005). The number of Sholl intersections was also greater for medial neurons than lateral neurons (**Figure 7G**; Two-way ANOVA, main effect of MD subnucleus; *F*(1, 315) = 84.98; *p* < 0.0001; Sidak’s multiple comparisons test: 105 − 165 μm from the soma, adjusted *p* values ≤ 0.005). Dendritic volume was greater in medial cells compared to lateral (**Figure 7D**; MD-M: 8562787 ± 2431041 µm^3^ (n = 7 neurons); MD-L: 4290298 ± 2248141 µm^3^ (n = 10 neurons); Mann-Whitney test; *U* = 5; *p* = 0.002). Finally, dendritic complexity (Pillai et al., 2012) was increased in medial neurons compared to lateral neurons (**Figure 7E**; MD-M: 245596 ± 119022 (n = 7 neurons); MD-L: 114497 ± 28860 arbitrary units (n = 10 neurons); Mann-Whitney test; *U* = 6; *p* = 0.0031).

**Figure 7:**
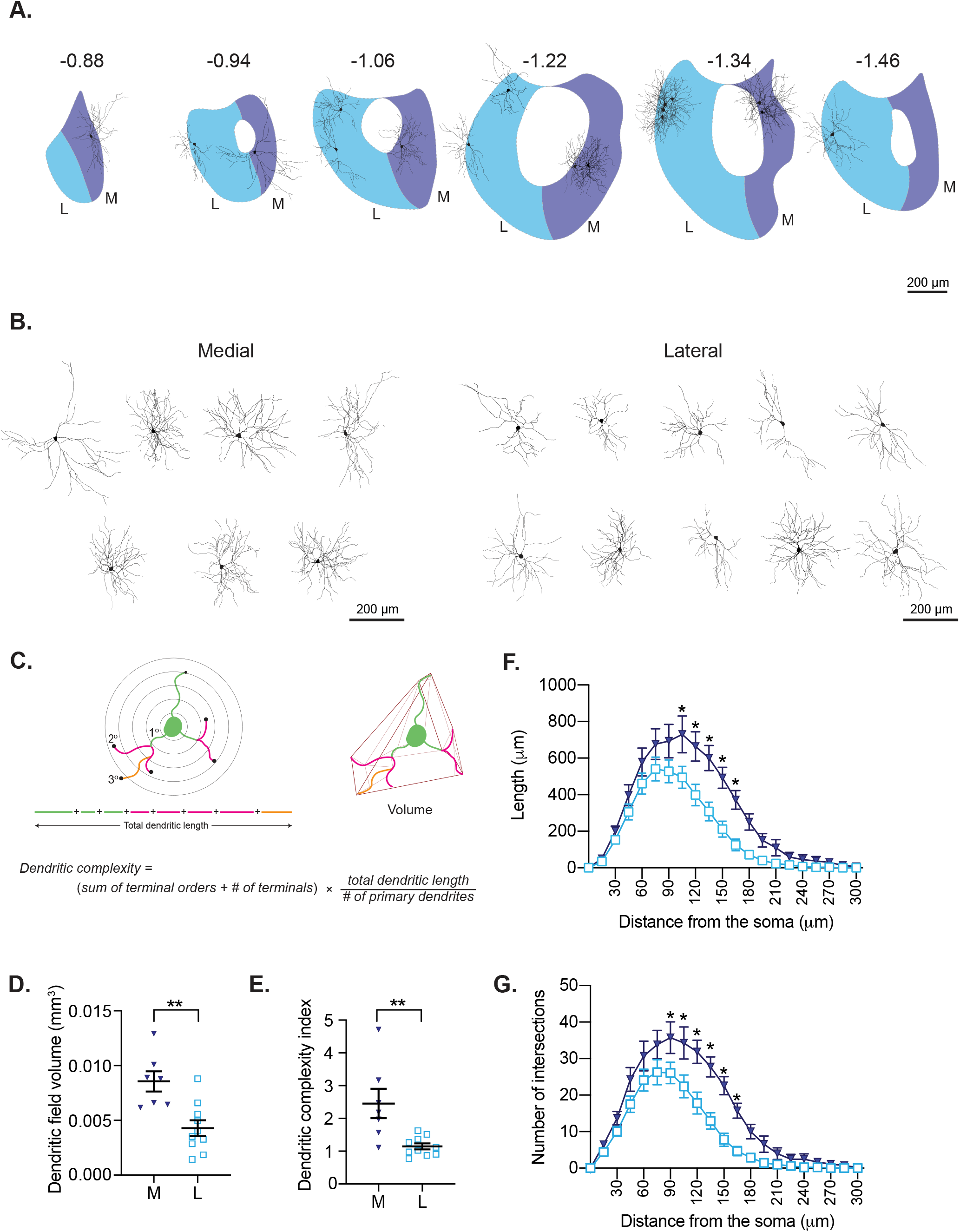
MD-M→mPFC neurons have larger, more complex dendrites than MD-L→mPFC neurons do. **A**. Map of approximate locations of reconstructed neurons. **B**. Tracings of medial and lateral neurons. **C**. Diagram example of analyses performed for reconstructed cells **D**. Medial neurons had greater field volume than lateral. **E**. Medial neurons had higher complexity index than lateral. **F. & G**. Sholl plots of dendritic length in each circle and total number of intersections with each circle. Scatter plots show all data points with the mean ± SEM. Significance is indicated at p<0.05 (*), p<0.01 (**). For **D, E**: two-tailed Mann-Whitney test. For **F, G**: Sidak’s test.

## DISCUSSION

### Summary of findings

Projections from MD→mPFC are required for executive functioning and social behavior (Parnaudeau et al., 2013, 2015; Bolkan et al., 2017; Schmitt et al., 2017; Ferguson and Gao, 2018; Rikhye et al., 2018). However, MD contains at least 3 identifiable subnuclei, two of which (MD-M and MD-L) provide substantial ascending input to mPFC. Given their spatial separation and distinct inputs and outputs, we hypothesized that the MD→mPFC projections in MD-M and MD-L are composed of populations of neurons with distinct intrinsic properties and morphologies. Here, we recorded from >100 neurons in the medial (MD-M) and lateral (MD-L) divisions of the mouse MD thalamus that were identified as projecting to ipsilateral medial prefrontal cortex (mPFC) using a retrograde tracer. Overall, we found that MD→mPFC neurons in the medial and lateral divisions of MD significantly differ in their physiological properties and dendritic morphology. MD-M cells had longer membrane time constants, higher cellular membrane resistance, more readily fired action potentials in response to small current inputs, and had larger, more complex dendritic arbors. MD-L cells on the other hand, tended to have faster membrane time constants, lower resistance, higher HCN-mediated voltage changes, and simpler dendritic arbors.

### Comparison to previous studies

This is the first study dedicated to understanding the physiology of adult MD→mPFC neurons. Surprisingly, only three published studies to our knowledge have measured whole cell physiology in MD neurons (Rajasekaran et al., 2007; Lee et al., 2011; Collins et al., 2018), none of which compared subnuclei. Collins *et al*. recorded primarily from MD-L→mPFC neurons in mice age P28 – P42. They reported RMP of around −65 mV (range −60 to −70 mV), R_m_ 175 MΩ (range 100 – 250), T_M_ in the low 20’s ms (range 15 – 35), an average of 3 rebound action potentials (range 0 – 7) in response to hyperpolarizing current steps, and fire up to about 60 Hz in response to +300 pA square pulses. There were no estimates of HCN channel activity reported. Lee *et al*. reported single cell recordings from MD neurons at P19 – P21. MD neurons had firing rates of around 60 Hz in response to +300 pA square pulses. The wildtype neurons in that report fired 1 – 6 rebound action potentials following membrane hyperpolarization.

The results we present differed from previous reports in the age of the animals (we recorded in 8 – 12 week old mice), in specifically comparing MD-M→mPFC neurons with MD-L→mPFC neurons, and in measuring and comparing HCN channel activity, one of the major contributors to membrane resistance. Our results complement estimates of rat MD-M and MD-L morphology that suggested a tendency toward larger neurons in MD-M compared to MD-L (e.g. (Kuramoto et al., 2017)). Our results also provide evidence in rodent that complement the histological features observed in nonhuman primates that form the basis for MD-M being dubbed “pars magnocellularis” (large cells) and MD-L known as “pars parvicellularis” (small cells) (Ray and Price, 1993).

### Strengths and limitations of this study

Strengths of the current study include the well-defined circuit (MD→mPFC neurons), the large number of neurons sampled, and multiple regions sampled along the DV / ML / and AP axes. Unfortunately, we had too few neurons reconstructed to be able to make meaningful correlations between neuron structure and function in this report. In this study, we did not distinguish between prelimbic and infralimbic cortices with our retrograde tracer injections. Both MD-M and MD-L project to mPFC, and the axon terminals extend over hundreds of micrometers and do not necessarily observe borders between anterior cingulate, prelimbic, and infralimbic cortex (Kuramoto et al., 2017). There may be a dorsal-ventral / lateral-medial topography of mPFC-MD connectivity in which ventral mPFC (**vmPFC**) (which is associated with monitoring of emotions) has stronger connectivity with **MD-M**. By contrast, dorsal mPFC (**dmPFC**), which is associated with monitoring actions and cognition, may have relatively increased connectivity with **MD-L** (Frith and Frith, 2006; Hunnicutt et al., 2014; Mátyás et al., 2014; Vertes et al., 2015; Alcaraz et al., 2016; Bolkan et al., 2017; Ährlund-Richter et al., 2019). Smaller, more precise injections that target either anterior cingulate, prelimbic, or infralimbic may further delineate cellular groupings. In addition, future work to determine molecular and genetic classifications of neurons through immunohistochemistry and single-cell RNA sequencing may help further delineate functional groups of projection neurons (Phillips et al., 2019).

### Implications of differences between MD-M and MD-L

MD-M→mPFC neurons have higher resistance and longer membrane time constants. We predict that this will provide MD-M→mPFC neurons a longer time window for temporal integration of synaptic inputs compared to MD-L→mPFC neurons. This could make MD-M neurons more sensitive to responding to poorly-coordinated inputs. This would be in contrast to MD-L cells that would potentially need inputs to be more closely-timed to summate and depolarize the membrane past the action potential threshold. Future work will involve understanding how different MD→mPFC neurons process incoming synaptic information. Even within MD-M and MD-L, there is a wide distribution of membrane resistance and time constants. The functional consequence of this is yet to be determined. Of note, variance of subthreshold intrinsic properties within thalamocortical neurons is hypothesized to improve information transfer in the circuit (Béhuret et al., 2015).

We observed that although MD-L→mPFC neurons have a higher rheobase (i.e. require more current to elicit spiking), their action potential threshold is more negative compared to MD-M→mPFC neurons. This implies that differences in voltage-gated sodium channels do not likely play a role in the decreased spiking input / output relationship for MD-L neurons compared to MD-M neurons. The relatively depolarized action potential threshold in MD-M neurons could possibly be explained by MD-M→mPFC neurons’ longer time constants, which can lead to deactivation of voltage-gated sodium channels in neurons (Howard and Rubel, 2010). This would need to be tested directly.

In prefrontal cortex and the CA1 region of hippocampus, principal cells can be distinguished based on HCN-channel mediated voltage sag (Jarsky et al., 2008; Dembrow et al., 2010; Maroso et al., 2016). Here, we did not find a specific cut-off between cell types based on voltage sag, but rather a smooth continuum of voltage sag that was on average higher in MD-L→mPFC neurons. There is growing interest in HCN channel activity for its role in axonal (Ko et al., 2016) and dendritic (Vaidya and Johnston, 2013) signaling, but also for its roles in epilepsy (Bernard et al., 2004), depression (Kim et al., 2017), neurodevelopmental disorders (Omrani et al., 2015; Brandalise et al., 2020), and neurodegenerative disorders (Chang et al., 2019). Further studies can parse out whether differences between cell types is due to expression levels, subunit composition, or biophysical properties.

Regarding differences in neuronal morphology, one implication of MD-M→mPFC neurons having larger and more complex dendritic arbors compared to MD-L→mPFC neurons is that each input onto the MD-M neuron may exert less influence over the neuron’s activity. This would be in contrast to MD-L neurons with their smaller dendritic arbors, thus fewer synaptic inputs would influence a given neuron’s input/output function. These morphological findings could complement the physiological findings to fit with a model of MD-M→mPFC neurons being optimized for sensitivity (more inputs that don’t need to be precisely timed) and MD-L→mPFC neurons optimized for specificity (fewer inputs that need to be more precisely timed to elicit action potentials).

Here, we identified and recorded from MD neurons that project to mPFC. MD neurons, however, are known to project to multiple downstream targets via branched axons (Kuramoto et al., 2017). Thus, the MD→mPFC neurons we recorded from may also project to other postsynaptic targets besides mPFC. Thus, the differences in intrinsic physiology we observed in MD→mPFC neurons in this report has implications for streams of information destined for mPFC and also for other postsynaptic targets of MD such as orbitofrontal cortex, the reticular nucleus of the thalamus, and the frontal eye fields.

Though differences in the prefrontal thalamocortical network have been identified in neuropsychiatric disorders, it is not yet known if differences in macroscopic network function reflect differences in the physiological properties of the neurons that make up the network. Understanding the physiology of the projections from MD→mPFC in typically developing animals as we have done here lays the foundation for understanding differences in the MD→mPFC network in neurodevelopmental and neuropsychiatric disorders.

## Supporting information

Statistics Table

## ACKNOWLEDGEMENTS

We thank Meredith McCarty, Aurora Weiden, Madelynn Campbell, Joy Adler, and Mendee Geist for their technical assistance. We thank members of the Brumback and Howard labs (especially Jessica Chancey and MacKenzie Howard) and Dan Johnston for helpful discussions.

## CONTRIBUTIONS

P.L. performed electrophysiological recordings. A.K. performed cellular morphology experiments, analyzed all cellular morphology data, and wrote portions of the paper. A.B. conceived of the project, designed all experiments, wrote all code, analyzed all data, and wrote the paper. All authors approved the final manuscript prior to publication.

